# Predicting real-life creativity using resting state electroencephalography

**DOI:** 10.1101/2023.07.28.550981

**Authors:** Fatima Chhade, Judie Tabbal, Véronique Paban, Manon Auffret, Mahmoud Hassan, Marc Vérin

## Abstract

Neuroscience research has shown that specific functional brain patterns can be related to creativity during multiple tasks but also at rest. Nevertheless, the electrophysiological correlates of a highly creative brain remain largely unexplored. This study aims to uncover resting-state networks related to real-life creativity using high-density electroencephalography (HD-EEG) and to test whether the strength of functional connectivity within these networks could predict individual creativity. We acquired resting-state HD-EEG data from 90 participants who completed a creativity questionnaire. We then employed connectome-based predictive modeling; a machine-learning technique that predicts behavioral measures from brain connectivity features. Using a support vector regression, our results revealed functional connectivity patterns related to high and low creativity in the gamma frequency band. In leave-one-out cross-validation, the combined model of high and low creativity networks predicted creativity scores with very good accuracy (r= 0.34, p= 0.0009). Furthermore, the model’s predictive power was established by an external validation on an independent dataset (N= 41), where we found a statistically significant relationship between the observed and predicted creativity scores (r= 0.37, p= 0.01). These findings reveal large-scale networks that could predict individual real-life creativity at rest, providing a crucial foundation for developing EEG network-based markers of creativity.

## INTRODUCTION

Higher order cognition is a range of complex and sophisticated thinking skills. Among the functions subsumed under this category is creativity; people’s ability to produce novel and valuable ideas. On account of the importance of creativity in cultural and technological progress and innovation, the neuroscience community showed a significant interest in studying its neural correlates in the brain. Over the past years, emerging evidence shows that complex brain functions like creativity are generated by large-scale networks of highly specialized and spatially segregated brain regions ^1^. Motivated by the enormous progress that has been made in developing neuroimaging tools, functional connectivity analysis measures the statistical relationship between neural activities of brain regions to investigate the interregional conversation that underlies creative mechanisms. Therefore, identifying functional brain networks related to creativity from neuroimaging data has emerged as a highly promising goal in neuroscience research.

Various neuroimaging data can be used to assess functional connectivity (FC), such as functional magnetic resonance imaging (fMRI), electroencephalography (EEG), or magneto electroencephalography (MEG). Using fMRI, FC studies have successfully demonstrated an association between brain activity patterns and higher-order cognitive traits such as fluid intelligence ^2^, attention ^3^, and memory ^4^. Importantly, considerable fMRI work explored the correlational relationships between patterns of FC and creative activity, during multiple tasks including divergent thinking, visual art production, figurative language production, musical improvisation, and poetry composition ^5^. Moreover, researchers have uncovered large-scale functional networks underlying creativity at rest ^5^, providing convincing evidence for the link between creativity and resting-state FC, defined as FC during a wakeful but resting state without performing goal-directed activities ^6^ ^7^ ^8^.

However, establishing a generalizable relationship between FC patterns and complex cognitive functions like creativity remains a challenge, given that the relationship is purely correlative in its nature. Correlation and regression analyses often fail to generalize their results to novel observations, mainly because of their tendency to overfit the data ^9^. Recently, to address this issue, cross-validation tests were introduced into FC analyses. Compared to correlation, cross-validation is a more powerful method to estimate a network-behavior relationship ^10^. Crucially, it facilitates testing the strength of the relationship in novel observation via both internal and external validation, permitting to establish the generalizability of the results to independent datasets. Consequently, a machine learning method with built-in cross-validation— connectome-based predictive modeling (CPM)—has been developed to extract the most relevant features from brain functional connectivity data in order to construct a predictive model of individual differences of cognitive behaviors ^9^. Interestingly, in a recent work, Beaty et al applied the CPM method and succeeded to identify a brain network associated with high creative thinking ability ^11^, using fMRI FC data acquired from participants during their engagement in the alternative uses task – a classic divergent thinking task– which measure participant’s ability to generate solutions to open problems ^12^.

In recent years, the recognized successes of FC research has inspired a growing interest in analyzing brain functional networks using EEG, as a practical, easy to use, and relatively low-cost neuroimaging technique. In contrast to fMRI, EEG reflects a direct measure of neural activity and allows to compute brain oscillations within specific frequency bands. These oscillations may reflect important properties of network interactions at local and large-scales ^13^. When combined with source reconstruction approaches ^1^, EEG can be used to study functional interactions among cortical regions. Several studies have used EEG to investigate functional patterns associated with creativity, during creative thinking ^14^ ^15^ ^16^ ^17^ or at rest ^18^. The studies have generally classified creativity based on creative tasks performance rather than individual differences in everyday and real-life creativity. Moreover, very few studies have used machine learning methods like CPM to explore how individual creativity influences functional connectivity during creative processing, and none, as far as we know, during the resting state.

In this work, we hypothesized the presence of an EEG resting-state neurophysiological marker of real-life creativity. Thus, we combined CPM with resting-state HD-EEG data acquired from 90 healthy adult participants who completed a questionnaire that assesses individual creative activities and achievements. The objective is to identify a large-scale functional network related to real-life creativity and to determine whether individual creativity can be reliably predicted based on the strength of functional connectivity within this network at rest. Furthermore, we conducted an external validation analysis to establish the predictive power of the resulting neural model in a second independent EEG data set of 41 healthy subjects.

## RESULTS

### Predictive Networks of Creativity

HD-EEG data was acquired from healthy adult participants during eyes-closed resting state. All participants completed a creativity questionnaire (Inventory of Creative Activities and Achievements questionnaire -ICAA-); a broad-based assessment of individual differences in real-life creativity. In this study, we aim to discover the existence of an EEG network that can predict ICAA creativity score. Thus, whole-brain functional networks were constructed for each participant by computing the source-space functional connectivity among 68 brain regions of interest, in five frequency bands (delta, theta, alpha, beta, and gamma). A correlation between all functional connections and creativity scores, followed by a statistical threshold (*p* < 0.01), was used to identify which functional edges are significantly related to creativity in each frequency band. To determine which of these networks have a predictive potential, we used the connectome-based predictive modeling approach (CPM) ^9^, where leave-one-out cross-validation (LOOCV) was performed to build and test network-based predictive models (i. e. internal validation). The analysis detected no relevant predictive networks of creativity in delta, theta, alpha, nor beta bands.

In the gamma band, it revealed a “high-creativity network” consisting of 8 edges positively correlated with ICAA scores, and a “low-creativity network” consisting of 26 edges negatively correlated with ICAA scores (total possible edges of 2,278). These predictive networks were consistent among 90% of all LOOCV folds. The high-creativity network exhibited dense functional connections among 6 regions, 3 within the default mode network (DMN), 2 within the visual association network, and 1 paracentral lobe structure (left). The low-creativity network showed diffuse connections among 20 regions across the whole brain. 6 were within the default network, 4 were within the sensorimotor network, 2 within the frontoparietal network, 2 within the ventral attention network, 1 within the visual network, and 5 temporal structures (**Figure 1**).

**Figure 1.**
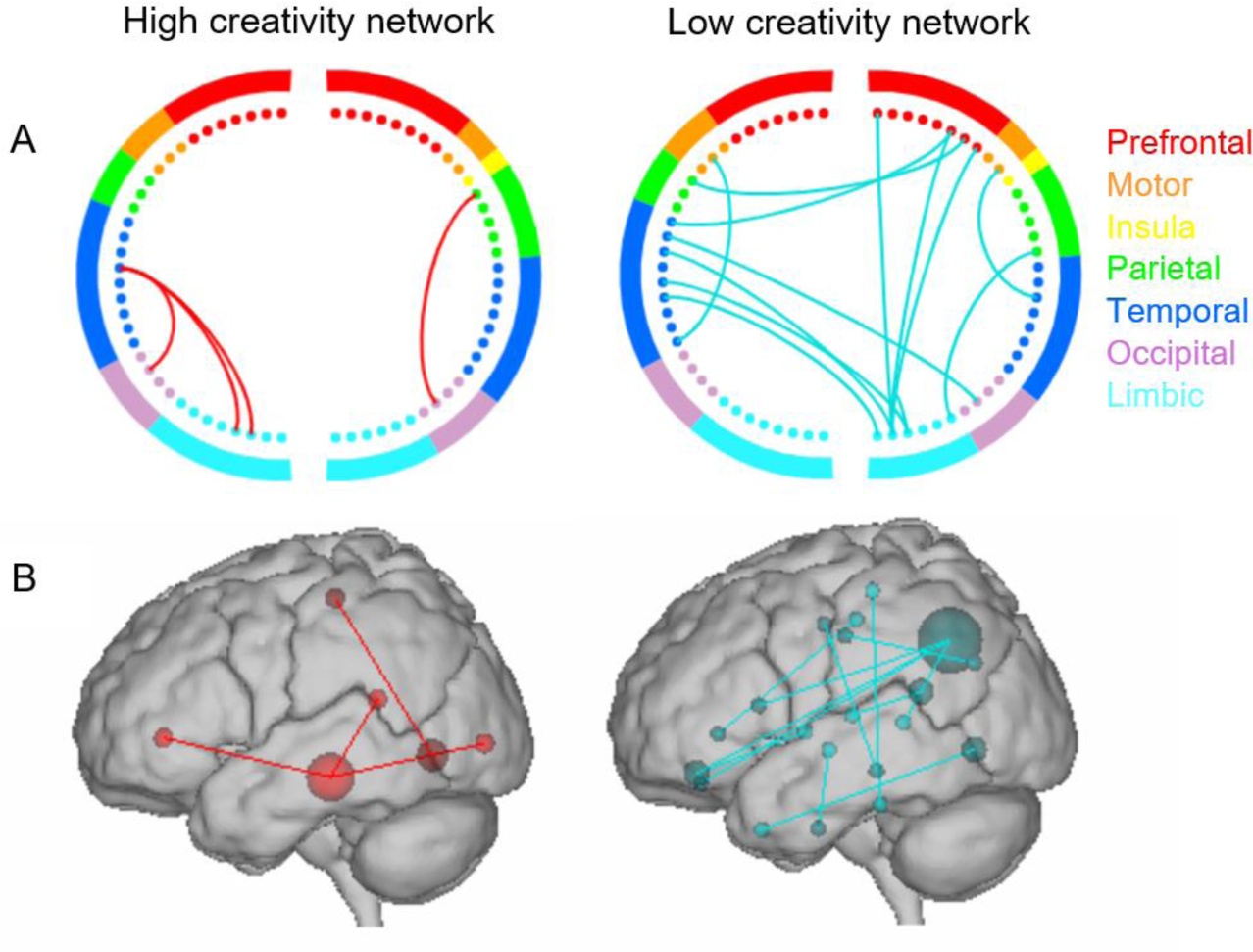
Depictions of high and low creativity networks. Circle plots (A) and glass brains (B). Colors within the circle plots correspond to brain lobes.

### Internal Validation: Prediction of Creativity from RS-EEG Data

As internal validation, we followed a LOOCV analysis (see figure 4). This step aims to test the predictive performance of the brain connectivity-based models that reflects the strength of functional connectivity within the high and the low creativity networks, respectively. In the leave-one-out loop, 90 rounds (i.e. number of participants) of cross-validation were performed, during which, one different participant was left out from building the model each time, then used to test the model performance. This caused slight differences in networks in each round, as well as the model and its performance. Thus, the final evaluation of the model was the average performance across the 90 folds.

We evaluated the predictive power of the model by assessing the statistical significance of the relationship between the observed and the model-predicted creativity scores. Results showed that high or low-creativity models alone could not robustly predict creativity. But interestingly, the combination of both networks could reliably predict real-life creativity (R=0.34, p-value=0.0009, MAE=38.4, R-squared=0.15) (**Figure 2**). Additionally, a non-parametric permutation test of ICAA scores (5000 times) was added to ensure that the obtained correlation is significantly better than expected by chance. The permutation test results confirmed the significance of the combined network (*p*-value = 0.02). This internal validation demonstrates that individual differences in real-life creativity could be predicted from the strength of RS functional connectivity within the combination of both high and low-creativity networks.

**Figure 2.**
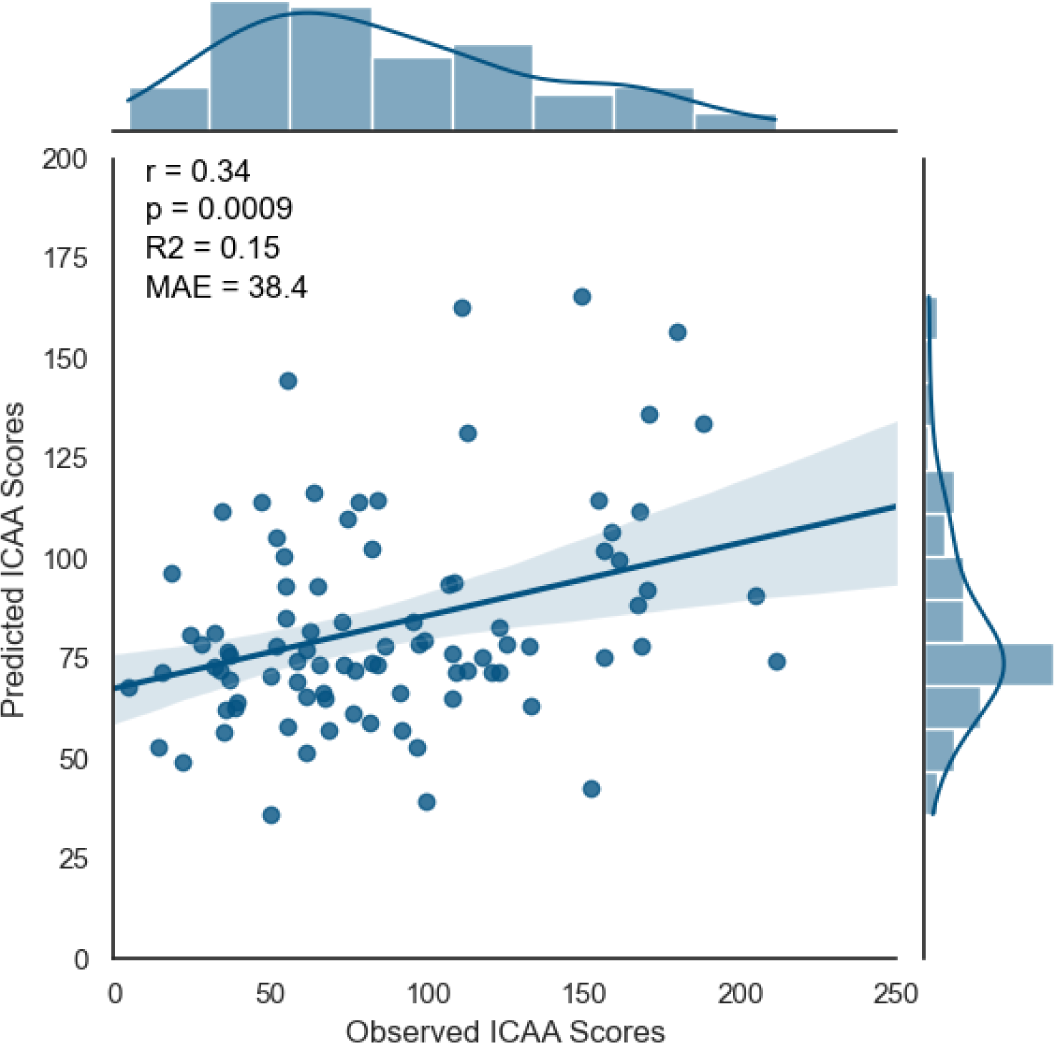
Relation between observed and predicted creativity scores (ICAA) in the internal validation. r = Correlation Coefficient, p = p-value, R2 = R-squared and MAE = Mean Absolute Error.

### External Validation: Prediction of Creativity Using Novel RS-EEG Data

Furthermore, to strengthen the generalizability of our predictive model, we conducted an external validation analysis using a second independent RS-EEG dataset.

High and low-creativity network strength values were thus computed for each subject in the new dataset. Then we used the trained model derived from the internal validation procedure to predict their creativity scores. Results revealed a significant predictive performance of the model (R=0.37, P value=0.01, MAE= 25.4, R-squared=0.14) (**Figure 3**). This external validation generalized the combined creativity model to a novel and distinct sample, indicating that participants with higher ICAA scores showed stronger functional connectivity within the high-creativity network and lower functional connectivity within the low-creativity network.

**Figure 3.**
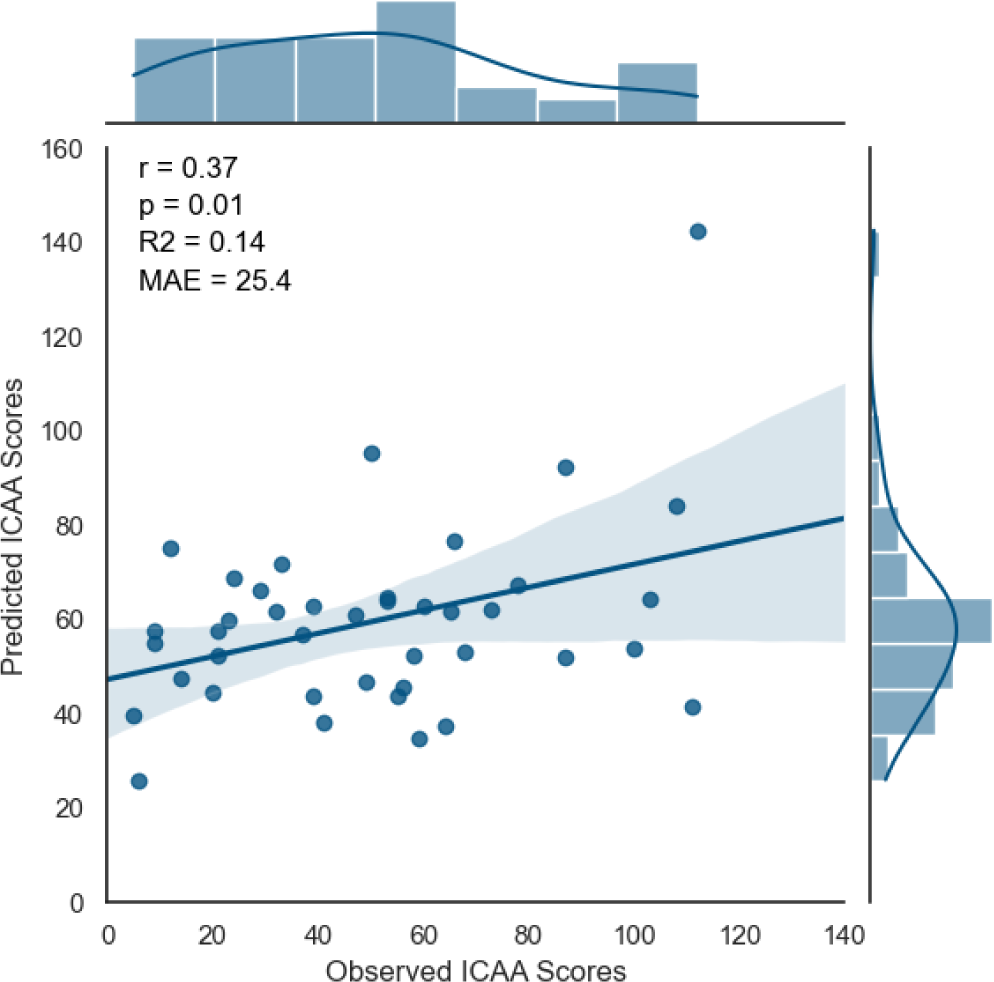
Relation between observed and predicted creativity scores (ICAA) in the external validation. r = Correlation Coefficient, p = p-value, R2 = R-squared and MAE = Mean Absolute Error.

### Confounder analysis

To explore potential confounding variation, we correlated each available co-variable (age, gender, education) with the creativity score (ICAA scores). Results showed no significant relation with neither age (*p*= 0.06), gender (*p*= 0.7) nor education (*p*= 0.36). Consequently, no cofounder effect was detected and thus considered as a variable in the predictive model.

## DISCUSSION

Using HD-EEG and a machine learning technique, we established a connectome-based model capable of predicting individual real-life creativity. The model is based on resting-state networks associated with high and low creativity, respectively. Critically, the model demonstrated its generalizability by successfully predicting creativity scores across an independent EEG dataset. These results indicate that creativity may be captured not only by task-induced variation of functional connectivity ^11^, but also by stable trait-level variation of intrinsic functional connectivity at rest. Several researches have examined resting-state functional connectivity (RSFC) and have revealed consistent RSFC large-scale networks in the human brain e.g. default mode network (DMN), visual network (VN), sensorimotor network (SMN), cognitive control network (CCN) ^19^ ^20^ ^21^ and salience network (SN) ^22^. Recent findings suggested that creativity can be associated with the strength of FC among specific patterns of interactions between brain regions of those resting-state networks. Our results are in line with these findings and increase evidence that real-life creativity can be predicted from an individual’s resting EEG connectivity profile.

The high-creativity network exhibits functional connections among core nodes of the DMN, which is involved in constructing dynamic mental simulations based on the past, the future, and imagination ^23^. Consistent with our findings, DMN was largely associated with creativity. The DMN activity has been positively correlated with higher creativity at rest, as divergent thinking ability ^24^ and verbal creativity ^25^. Moreover, DMN regions are often involved in task-induced creativity such as visual creativity ^26^, creative story generation ^27^, and insightful problem solving ^28^. In addition to the default mode, the high creativity network included core regions of the visual association network. Visual association is necessary to identify objects and interpret the same object across different specific contexts ^29^. Evidence showed that visual association is closely associated with memory performance ^30^ ^31^ ^32^ ^33^. Recent work suggested an important role for visual association areas in associative memory ^34^ —the ability to bind together information that was previously unrelated —underlies the formation of episodic memories ^35^. The engagement of DMN and associative memory may be in line with a theory of episodic memory; the *constructive episodic simulation hypothesis.* The theory postulates that memory and imagination, together, implicate flexible recombination of episodic memories such as places, people, objects, and details of events. This could be very interesting since, in part, creativity is supported by our abilities to recall memories and envision the future, especially at rest. This flexible nature of episodic processes seems to be particularly involved in creativity, which requires connecting both memory and imagination in new, original and meaningful ways. This builds on other works that have indicated that episodic memory is closely related to creativity ^36^ ^37^.

The low-creativity network, on the other hand, exhibits diffuse functional connections among several resting state networks, especially DMN and sensorimotor network, in addition to temporal structures. Although DMN regions are associated with high creative ability, default activity was also linked with prepotent response tendencies ^38^ and strong semantic associations ^39^. Additionally, sensorimotor regions were found to support procedural previously learned information ^40^. This could suggest that low creativity may be characterized by increased interactions among regions that support memory and automatic common associations and representations that are in turn, not effectively regulated by high-creativity brain regions.

In our results, neither high creativity nor low creativity network alone could predict robustly individual creativity. But interestingly, together, they construct an effective predictive model. A possible explanation is that, at rest, the more people activate the high creative network that supports flexible reconstruction and recombination of information, and at the same time, the less they activate the low creative network that supports common and automatic information, the more creative they are in their real-life activities and achievements.

Our statistical analysis showed that only the gamma frequency band highlighted a robust internal and external predictive network of high real-life creativity, and no other significant outcomes were found in other EEG frequency bands. These observations may be explained by the fact that among all frequencies, the gamma band (30–100 Hz) in EEG is most closely associated with high-order cognitive function ^41^. It was shown to reflect multiple cognitive processes such as attention, language, binding and object representation ^42^. Interestingly, rapid oscillatory activity in the gamma band has been identified as a fundamental mechanism for effectively encoding and retrieving episodic memories ^43^ ^44^. Furthermore, it was suggested that gamma band activity is involved in creative thinking ^45^, and plays a critical role in creative insights, as was demonstrated by a study where EEG recordings revealed a burst of gamma activity beginning shortly prior to insightful solutions for verbal problems ^28^. However, when examining neural mechanisms underlying creativity using EEG, the gamma band wasn’t the only frequency band that highlighted creative ability. Alpha band activity on the other hand was reported to correlate with divergent thinking ability ^46^ ^47^ ^48^ ^49^. While there is a general assumption of the involvement of alpha band activity in divergent thinking processes, inconsistent results have been reported for alpha and beta bands. This inconsistency can be attributed to the heterogeneity of employed methods and participant’s samples as well as the relatively limited number of studies investigating EEG correlates of creativity.

Moreover, numerous functional connectivity studies have examined the statistical associations (i.e. correlations) between creativity and brain connectivity. Some of these studies have misinterpreted these associations as predictions. Yet, significant correlation within a sample does not necessarily indicate predictive ability, as discussed in ^50^. Thus, the advantage of this work lies in its use of cross-validation to rigorously assess the presence of a resting state FC-creativity relationship. By evaluating the predictive power of creativity networks, this approach provides a robust test of this relationship. Generally, there are two approaches of cross-validation, (1) a k-fold cross-validation in which the data are split into k different subsets or folds, and (2) leave one out cross-validation (LOOCV), which was performed in our analysis.

LOOCV is the simplest form and the most popular choice. Many researchers suggested using the k-fold method rather than LOOCV, considering that it gives less variable estimates of the prediction error than those from LOOCV ^51^. On that account, we also tested the k-fold approach and compared the outcomes. The results showed no consistency between the two methods and significantly better performance of LOOCV in our data. Nevertheless, the robustness of our LOOCV-derived model was largely due to the complementary external validation that allowed our experimental design to be particularly powerful and our model to be generalized to an independent dataset.

Another consideration in this study is that we used a self-reported questionnaire —the ICAA questionnaire — to assess participants’ creativity. The questionnaire provides scales for the frequency of engagement in everyday creative activity and the level of creative achievement across the most common eight creative domains. It offers a broad assessment and a quick gathering of a large amount of information on participants’ real-life creative behavior across multiple domains and levels. We ensured that participants completed the questionnaire in private with the assurance of confidentiality and anonymity to promote more truthful and accurate responses. Nonetheless, like all self-reported data collection, they are subject to limitations of honesty and reliability, given that people tend to be consciously or unconsciously biased when they report on their own experiences. However, the reliability and validity of the ICAA scores have been supported by a rigorous formal test analysis conducted on a large sample size of 1556 individuals, as demonstrated in ^52^.

In the end, it is crucial to acknowledge that studying creativity poses a major challenge due to its multifaceted and complex nature, which manifests in various forms and engages multiple brain regions. Unlike other aspects of cognition that have been attributed to localized brain activity, creativity is a network phenomenon that involves complex neural interplay between multiple brain regions across the whole brain. Considerable further research is needed to uncover the diverse manifestations of creativity and the distinct roles of different networks in the creative brain. Network-based approaches, such as connectome-based modeling, provide promising tools to address these questions. However, it is important that future researchers continue to further explore the large-scale networks and network dynamics underlying creative processes in different task contexts, such as divergent thinking and creative arts. EEG-HD provide a great opportunity to analyze functional network dynamics with high temporal precision, due to its excellent temporal resolution. An important advantage especially when studying FC of complex cognitive processes.

## METHODS

### Participants

Two independent datasets have been used for the study.

Dataset 1 was collected as part of the current study. A total of 98 healthy participants were recruited at the University Hospital Centre of Rennes from local and surrounding communities (60 female, mean age= 39.6 y ± SD= 12.7). To ensure a diverse population, participants from various creative domains such as art, dance, music, and sciences were selected. All participants were French native speakers, aged between 18 and 68 years, and with normal or corrected vision. They reported no situation or history of cognitive disability, neurological disorders, or medication that can affect the central nervous system. The study was approved by the ethical committee of the University Hospital Centre of Rennes (agreement n°20-171) and each participant provided written informed consent.

Dataset 2 was part of a different study at the University of Marseille in France. It consists of 52 healthy participants (28 female, mean age= 45.8 y ± SD= 17.3). The same inclusion criteria were applied. The study was approved by the “Comité de Protection des Personnes Sud Méditerranée” (agreement n°10-40) and each participant provided written informed consent prior to acquisition.

### Behavioral Assessment

Creativity was assessed by the Inventory of Creative Activities and Achievements questionnaire (ICAA): a broad-based assessment of individual differences in real-life creativity ^53^. The questionnaire provides independent scales for the frequency of engagement in everyday creative activity and the level of creative achievement across 8 creative domains (i.e., literature, music, art and crafts, creative cooking, sport, visual arts, performing arts, and science and engineering). For each participant, two scores were calculated: the creative activity score (Cact) based on the frequency of engagement in everyday creative activity, and the creative achievement score (Cash) based on the level of publicly acknowledged creative achievement. In this study, we used the total creativity score by summing Cact and Cash to obtain one score for each participant (C total). C total score can range between 1 (for least creative) and 472 (for most creative).

### EEG Data Acquisition

EEG recordings were collected during a resting state. During the acquisition session, participants were seated comfortably in a dimly lit room and asked to rest and relax for 5 to 6 minutes while closing their eyes without falling asleep. In Dataset 1, the EEG was acquired with the high-density 256-channel HydroCel Geodesic Sensor Net (Electrical Geodesics Inc., Eugene, OR, United States) at a sampling rate of 1000 Hz. We referenced all electrodes to Cz, and impedance remained below 50 kΩ. In Dataset 2, EEG recordings were collected using a 64-channel Biosemi ActiveTwo system at a sampling rate of 2048Hz.

### EEG Data Preprocessing

We applied a semi-automated preprocessing protocol for both datasets. First, for each participant, 5-6 minutes signal was segmented into non-overlapping 40 seconds epochs. The epochs of Dataset 2 were resampled at 1000 Hz to equalize the sampling frequency. Then, we used an automatic protocol on AUTOMAGIC®; an open-source MATLAB-based toolbox for EEG preprocessing ^54^. The protocol consisted of three main steps: i) bad channel identification, ii) artifact correction including a pass filter between 1 and 45 Hz and EOG regression, and iii) interpolation of detected bad channels using neighboring electrodes within a 4-5 cm radius. Furthermore, to ensure good signal quality, each epoch was visually inspected, and additional bad channel detection and interpolation were performed if needed. After interpolating, we applied a 15% threshold, meaning the exclusion of every epoch that needed more than 15% of electrodes to be interpolated. Finally, we selected three artifact-free epochs of 40 seconds in length for each participant. Due to their poor signal quality, 8 participants were excluded from the first dataset and 11 from the second.

### Brain Network Construction

Functional brain networks were estimated using the ‘EEG source connectivity’ method ^1^, combined with a sliding window approach as detailed in ^55^. The method includes two main steps: i) solving the EEG inverse problem to estimate the cortical sources in order to reconstruct their temporal dynamics and ii) measuring the functional connectivity between the reconstructed scout time series.

#### Source estimation

In order to solve the inverse problem, the weighted minimum norm estimate (wMNE) algorithm ^56^ was used to estimate regional time series between predefined regions of interest (ROIs). To define ROIs, we used the Desikan Killiany atlas, which parcellate the cortical surface into 68 ROIs ^57^.

#### Functional connectivity

The functional connectivity between the 68 regional time series was obtained using the phase-locking value metric (PLV). The combination of (wMNE/PLV) proved its efficiency to identify the cortical brain networks from scalp EEG recordings at rest ^58^, and during cognitive activity ^59^ ^60^.

We filtered the reconstructed regional time series in different frequency bands (delta: 1–4 Hz; theta: 4–8 Hz; alpha: 8–13 Hz; beta: 13–30 Hz and gamma: 30-45 Hz) and we applied the sliding window method in which PLV was calculated over its data points. Then PLV was averaged across sliding windows. As a result, we obtained one dynamic PLV and one static PLV for each frequency band, and for each participant.

### Connectome-based Predictive Modeling

We employed connectome-based predictive modeling (CPM) ^61^ on the first dataset to construct predictive networks that can be used to estimate individual real-life creativity (C total) from resting-state EEG functional connectivity. CPM is a recently developed method for identifying and modeling functional brain connections related to a behavior of interest; real-life creativity in our case. The connectome network is then used to predict the behavior of novel participants whose data were not used in model creation. The method was previously employed and described in a number of studies that showed its results in predicting cognitive variables such as attention ^3^ and fluid intelligence ^2^. As well as to predict network alteration in a number of brain disorders such as sleep disorders ^62^ and anxiety ^63^.

For a detailed description of CPM, refer to Shen et al. ^9^. Hereafter, a brief summary of the processing pipeline including both training and testing procedures (**Figure 4**).

**Figure 4.**
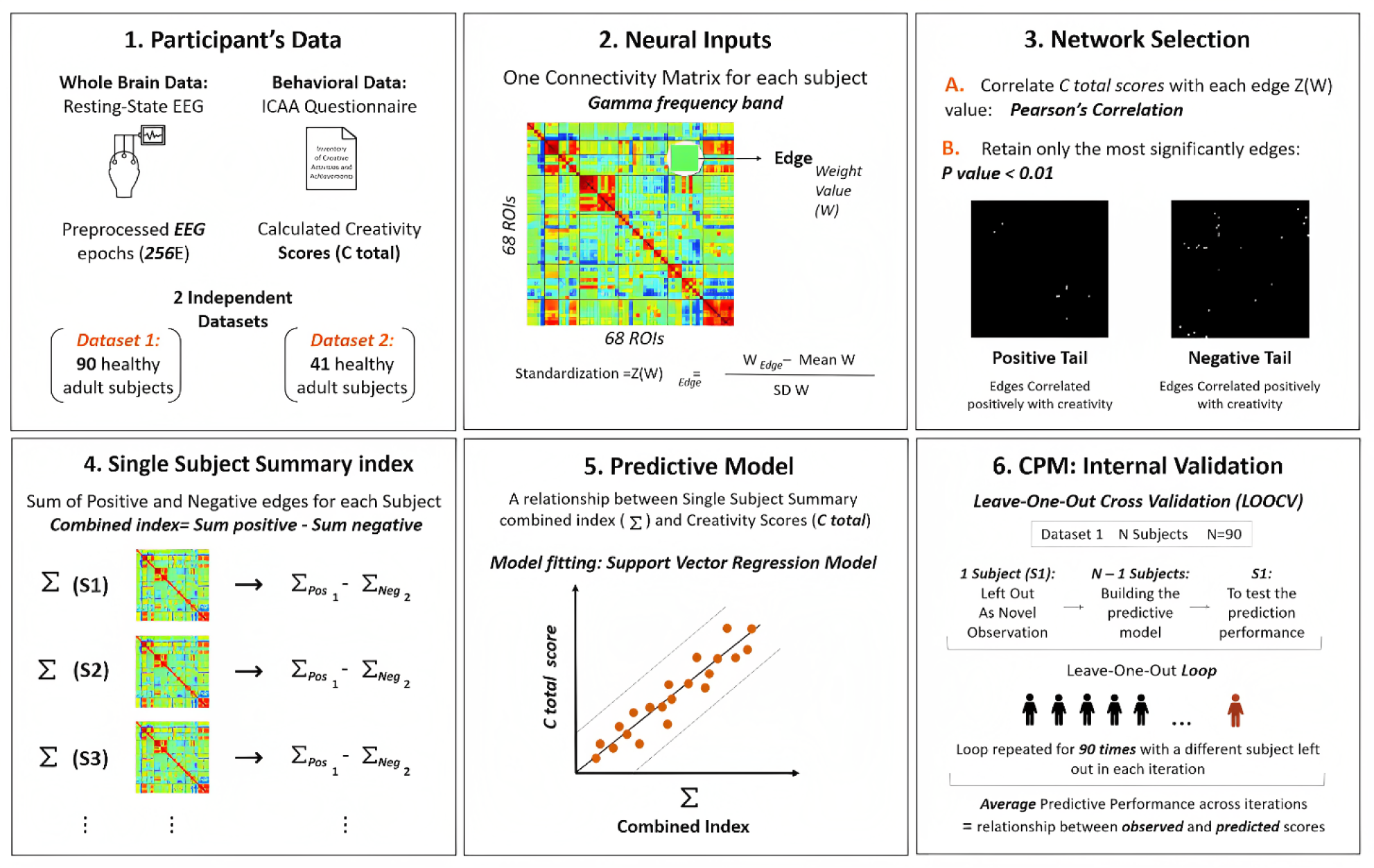
A full pipeline of the employed CPM method as described in the Connectome-based Predictive Modeling section.

#### Prepare the model inputs

i) The vector of behavioral values represented by creativity scores (C total) for each participant and ii) the standardized weight values of each edge in the functional connectivity matrix of each participant. For edge standardization, as used in multiple previous studies, a Z-transformation was performed on each edge by calculating the difference between its weight and the mean weight divided by the standard deviation across the subjects in the training set ^64^ ^65^. The same parameters acquired during the training procedure (mean and standard deviation) were used to standardize the connectivity edges of the testing set.

#### Identify the predictive edges

Using Pearson’s correlation, each edge (i.e., standardized weight value) in the connectivity matrix was correlated with creativity scores, while applying a threshold (P < 0.01) in order to retain the most significant positively and negatively correlated edges. This step resulted in two reconstructed networks: a high-creativity network (i.e., edges correlated positively with C total scores) and a low-creativity network (i.e., edges correlated negatively with C total scores).

#### Build the predictive model

The edge strengths were computed in both positive and negative tails of correlation. By combining both creativity networks, we calculated the summed index. This latter was obtained by summing standardized weight values of all connections of the positive tail and subtracting those of the negative tail. Then, a support vector regression (SVR) model was employed to respectively relate behavioral and connectivity strength values of the training set. SVR aims to find the hyperplane that maximizes the margin between predicted and actual score values. It can handle high dimensional data advantageously and provides a non-linear model that can effectively capture complex relationships in human brain data ^66^ ^67^. Here, we used the radial basis function (RBF) as a non-linear kernel.

#### Test the predictive model

The trained model was used to predict the creativity scores of the testing participant. Leave-one-out cross-validation (LOOCV) was applied. This strategy consists of removing one subject from the data as a novel observation and using N-1 subjects to build the predictive model, then using the novel subject to test its prediction performance. This step is repeated 90 times (i.e. number of participants) with a different subject left out in each iteration. The resulting performance presents the average performance across all iterations.

#### Assess the predictive model

To assess the predictive power of the established model, we evaluated the relationship between the observed behavior score and the predicted behavior score using several metrics: Pearson’s correlation (R), parametric p-value, mean absolute error (MAE), and the coefficient of determination (R-squared).

Additionally, to assess the statistical significance of the prediction results for creativity networks, we performed a non-parametric permutation test by randomly shuffling the creativity scores of all participants 5000 times. In each permutation, we randomly assigned one individual’s score to another individual’s connectivity data, repeated the leave-one-out cross-validation procedure, and calculated the correlation measure averaged across folds. The 5000 correlation coefficients generated a null distribution of R-values. Then, the non-parametric p-value was calculated as the number of permutations where the prediction correlation is greater or equal to the true prediction correlation divided by the total number of permutations.

### External Validation

Due to the slight differences in the resulting predictive networks in each iteration within the leave-one-out loop, we defined the “final” positive and negative predictive edges as those that were persistent in at least 90% of iterations. These positive and negative predictive networks derived from the internal validation (dataset 1) were then applied to a second independent dataset (dataset 2) for external validation. Incorporating an independent dataset is highly advised to establish the generalizability of the CPM model ^68^ ^69^.

First, edge z-transformation was applied using the same standardization parameters obtained from Dataset 1. Then, FC strength was computed within high and low-creativity networks, and the trained SVR models were applied to predict the creativity score for each participant in the new sample. Here, the same assessment metrics were used to evaluate the predictive models performance (R, p-value, MAE and R-squared).

### Confounder Analysis

To explore potential confounding variation, we correlated each co-variable with the creativity score (i.e. the dependent variable) using Pearson’s correlation for age and education, and Point-Biserial Correlation for gender. Education level was measured based on the official French education system, from 0 (without any diploma) to 5 (diploma of level BAC+5 or more).

## Data availability

The datasets generated during the current study are presents in the paper. Additional details are available from the corresponding author on request.

## Code availability

All MATLAB codes related to connectome-based predictive modeling is available online at https://www.nitrc.org/projects/bioimagesuite/, as well as the visualization software (https://www.nitrc.org/projects/bioimagesuite/).

## Acknowledgments

This work was partly funded by a Research Grant from the doctoral school of biology and health at the University of Rennes. We thank CECAP association « Comité d’entente et de coordination des associations de parkinsoniens » for partly supporting the study. We are also thankful to Rennes University Hospital Center for their High-density electroencephalogram acquisition facility. We are grateful to Ph.D. student Sahar Yassine for her technical advice.

## Author contributions

Supervision and funding acquisition: M. H. and M. V., experimental design: F. C., M. H., M. V. and V. P., data collection; F. C. (dataset 1) and V. P. (dataset 2), data preprocessing: F. C., methodological design: F. C., J. T., M. H., and M. V., algorithm coding: J. T., formal behavioral and network analysis: F. C., investigation and visualization: F. C., writing – original draft: F. C., writing – review and editing: all authors.

